# Immunoinformatics Prediction of Epitope Based Peptide Vaccine Against Mycobacterium Tuberculosis PPE65 family Protein

**DOI:** 10.1101/755983

**Authors:** Mustafa Elhag, Anfal Osama Mohamed Sati, Moaaz Mohammed Saadaldin, Mohammed A. Hassan

## Abstract

**Introduction:** Tuberculosis (TB) is a serious disease with varying rates of mortality and morbidity among infected individuals which estimates for approximately two million deaths/year. The number of deaths could increase by 60% if left untreated. It mainly affects immune-compromised individuals and people of third world, due to poverty, low health standards, and inadequate medical care. It has varying range of manifestations that is affected by the host immune system response, the strain causing the infection, its virulence, and transmissibility.

**Materials and methods:** A total of 1750 *Mycobacterium Tuberculosis* PPE65 family protein strains were retrieved from National Center for Biotechnology Information (NCBI) database on March 2019 and several tools were used for the analysis of the T- and B-cell peptides and homology modelling.

**Results and conclusion:** Four strong epitope candidates had been predicted in this study for having good binding affinity to HLA alleles, good global population coverage percentages. These peptides are YAGPGSGPM, AELDASVAM, GRAFNNFAAPRYGFK and a single B-cell peptide YAGP.

This study uses immunoinformatics approach for the design of peptide based vaccines for M. tuberculosis. Peptide based vaccines are safer, more stable and less hazardous/allergenic when compared to conventional vaccines. In addition, peptide vaccines are less labouring, time consuming and cost efficient. The only weakness is the need to introduce an adjuvant to increase immunogenic stimulation of the vaccine recipient.

## INTRODUCTION

Tuberculosis (TB) is a serious disease with varying rates of mortality and morbidity among infected individuals which estimates for approximately two million deaths/year. The number of deaths could increase by 60% if left untreated. [1][2][3] It mainly affects immune-compromised individuals and people of third world, due to poverty, low health standards, and inadequate medical care. [4][5][6] It has varying range of manifestations that’s affected by the host immune system response, the strain causing the infection, its virulence, and transmissibility. [7][8] These manifestations include but are not limited to: Constitutional symptoms for all forms of TB include fever, night sweats, and failure to thrive (children) or weight loss (adults). [9] Milliary tuberculosis, skeletal deformities, central nervous system involvement (tuberculosis meningitis TBM) and other organs involvement i.e., cervical lymphadenitis. [10][11][12][13]The main site of infection which follows the inhalation of aerosols or droplets containing *Mycobacterium Tuberculosis* is the alveoli in lungs where they enter mononuclear cells and interact with them following phagocytosis, causing damage, notably necrosis, cavitation, and coughing of blood. [5][14][15][16][17]

*Mycobacterium Tuberculosis* is a distinctive Acid fast bacilli that is obligate pathogens. [18]They differ from other bacteria in that a very large portion of its coding region is devoted to the production of lipogenesis and lipolysis enzymes, and to two new families of glycine-rich proteins with a repetitive structure that may represent a source of antigenic variation. [19]*M. tuberculosis* contains around 4,000 genes, one of which is the Proline Proline Glutamatic acid PPE gene, which comes from PE/PPE/PE-PGRS gene family which is present exclusively in genus Mycobacterium, accounts for approximately 10% of the M. tuberculosis genome and plays a role in the makeup of cell surface markers thus, helping in genetic variations, adhesion and invasion of host defense cells. [20][21][22][23][24]Studies suggest that variations in these genes may underlie the basis of virulence attenuation. [24][25]

One of the *M. tuberculosis* PPE proteins, the PPE65 (*Rv3621c*) harbored in the Region of Difference (RD8) is suggested to be potential virulence determinant.[24] It’s found to be overexpressed in macrophages infected with various clinical isolates of *M. tuberculosis* and it’s particularly, a good B-cell antigen with a higher antibody response.[23][26][27][28] PPE65 is thought to be a suppressor of pro-inflammatory cytokines such as TNF-α and IL-6 while it also induces high expression of anti-inflammatory IL-10 co-operated by PE32 protein causing inhibition of T-helper 1 cells. [29]

Morbidity and mortality rates of TB steadily dropped due to, improved public health practices and widespread use of the Bacille Calmette–Guérin (BCG) vaccine, as well as the development of antibiotics. [7] Then numbers of new cases started increasing. Hence, homelessness and poverty increased and the AIDS emerged, with its destruction of the cell-mediated immune response in co-infected persons. [12][30][31][32]This global crisis was followed by the emergence of multidrug resistance and extensively drug resistant strains in countries like the former Soviet Union, South Africa, and India, where some antibiotics are available in lower quality or are not used for a sufficient time to control the disease according to recommended regimens [5][33][34][35][36][37][38][39][40]In addition, BCG, the widely administered live attenuated vaccine against TB, is inconsistently effective in adults.[10][38][41][42]Therefore, development of new specific vaccine preventive against *M. tuberculosis*, is becoming a major priority of *M. tuberculosis* investigators.

This study aims to suggest new possible peptides for a peptide-based vaccine for *M. tuberculosis*, targeting PPE65 protein as an immunogen using an immunoinformatics approach. Reverse vaccinology (peptide based vaccines) is becoming more favoured over conventional vaccine methodologies. This is owes to the fact that reverse vaccine techniques are much less costing, more time and labour saving and makes patients less prone to hazards.[43][44][45] peptide vaccines are based on the chemical approach to synthesize the computationally predicted suitable B-cell and T-cell epitopes that are immunodominant and can induce specific immune responses.[43]this study differs from other studies for being the first study to use PPE65 protein as an immunogenic target for a reverse vaccine-design based study.

## MATERIALS AND METHODS

### Protein Sequence Retrieval

A total of 1750 *Mycobacterium Tuberculosis* PPE65 family protein strains were retrieved from National Center for Biotechnology Information (NCBI) database on March 2019 in FASTA format. These strains were submitted from different parts of the world and we collected them for immunoinformatics analysis. The retrieved protein strains had length of 284 base pairs with name “PPE65 family protein”

### Determination of conserved regions

The retrieved sequences of *Mycobacterium Tuberculosis* PPE65 family protein were subjected to multiple sequence alignment (MSA) using ClustalW tool of BioEdit Sequence Alignment Editor Software version 7.2.5to determine the conserved regions. Molecular weight and amino acid composition of the protein were also obtained using this tool.[46][47]

### Sequenced-Based Method

The reference sequence (WP_096833455.1) of *Mycobacterium Tuberculosis* PPE65 family protein was submitted to different prediction tools at the Immune Epitope Database (IEDB) analysis resource (http://www.iedb.org/) to predict various B and T cell epitopes. Conserved epitopes would be considered as candidate epitopes for B and T cells and were subjected for further analysis.[48]

### B-Cell Epitope Prediction

B cell epitopes is the portion of the vaccine that interacts with B lymphocytes. Candidate epitopes were analysed using several B cell prediction methods from IEDB (http://tools.iedb.org/bcell/), to identify the surface accessibility, antigenicity and hydrophilicity with the aid of random forest algorithm, a form of unsupervised learning. The Bepipred Linear Epitope Prediction 2was used to predict linear B cell epitope with default threshold value 0.533 (http://tools.iedb.org/bcell/result/). The Emini Surface Accessibility Prediction tool was used to detect the surface accessibility with default threshold value 1.000 (http://tools.iedb.org/bcell/result/). The Kolaskar and Tongaonker Antigenicity method was used to identify the antigenicity sites of candidate epitope with default threshold value 1.032 (http://tools.iedb.org/bcell/result/). The Parker Hydrophilicity Prediction tool was used to identify the hydrophilic, accessible, or mobile regions with default threshold value 1.695.[49–53]

### T- Cell Epitope Prediction MHC Class I Binding

T cell epitopes is the portion of the vaccine that interacts with T lymphocytes. Analysis of peptide binding to the MHC (Major Histocompatibility complex) class I molecule was assessed by the IEDB MHC I prediction tool (http://tools.iedb.org/mhci/) to predict cytotoxic T cell epitopes. The presentation of peptide complex to T lymphocyte undergoes several steps. Artificial Neural Network (ANN) 4.0 prediction method was used to predict the binding affinity. Before the prediction, all human allele lengths were selected and set to 9amino acids. The half-maximal inhibitory concentration (IC50) value required for all conserved epitopes to bind at score less than 100 were selected.[43, 54–59]

### T- Cell Epitope Prediction MHC Class II Binding

Prediction of T cell epitopes interacting with MHC Class II was assessed by the IEDB MHC II prediction tool (http://tools.iedb.org/mhcii/) for helper T cells. Human allele references set were used to determine the interaction potentials of T cell epitopes and MHC Class II allele (HLA DR, DP and DQ). NN-align method was used to predict the binding affinity. IC50 values at score less than 500 were selected.[60–63]

### Population Coverage

In IEDB, the population coverage link was selected to analyse the epitopes. This tool calculates the fraction of individuals predicted to respond to a given set of epitopes with known MHC restrictions (http://tools.iedb.org/population/iedbinput). The appropriate checkbox for calculation was checked based on MHC-I, MHC-II separately and combination of both.[64]

### Homology Modelling

The 3D structure requires PDB ID which was obtained using raptorX (http://raptorx.uchicago.edu) i.e. a protein structure prediction server developed by Xu group, excelling at predicting 3D structures for protein sequences without close homologs in the Protein Data Bank (PDB). USCF chimera (version 1.13.1rc) was the program used for visualization and analysis of molecular structure of the promising epitopes (http://www.cgl.uscf.edu/chimera).[65, 66]

## RESULTS

### Multiple Sequence Alignment

The conserved regions and amino acid composition for the reference sequence of *Mycobacterium Tuberculosis* PPE65 family protein are illustrated in figure 1 and 2 respectively. Glycine and alanine were the most frequent amino acids (Table 1).

**Table 1:** Molecular weight and amino acid frequency distribution of the protein

| Amino Acid | Number | Mol% |
| --- | --- | --- |
| Ala | 103 | 24.94 |
| Cys | 0 | 0 |
| Asp | 12 | 2.91 |
| Glu | 13 | 3.15 |
| Phe | 15 | 3.63 |
| Gly | 47 | 11.38 |
| His | 3 | 0.73 |
| Ile | 9 | 2.18 |
| Lys | 9 | 2.18 |
| Leu | 35 | 8.47 |
| Met | 11 | 2.66 |
| Asn | 11 | 2.66 |
| Pro | 28 | 6.78 |
| Gln | 19 | 4.6 |
| Arg | 5 | 1.21 |
| Ser | 29 | 7.02 |
| Thr | 25 | 6.05 |
| Val | 22 | 5.33 |
| Trp | 9 | 2.18 |
| Tyr | 8 | 1.94 |

**Figure 1:**
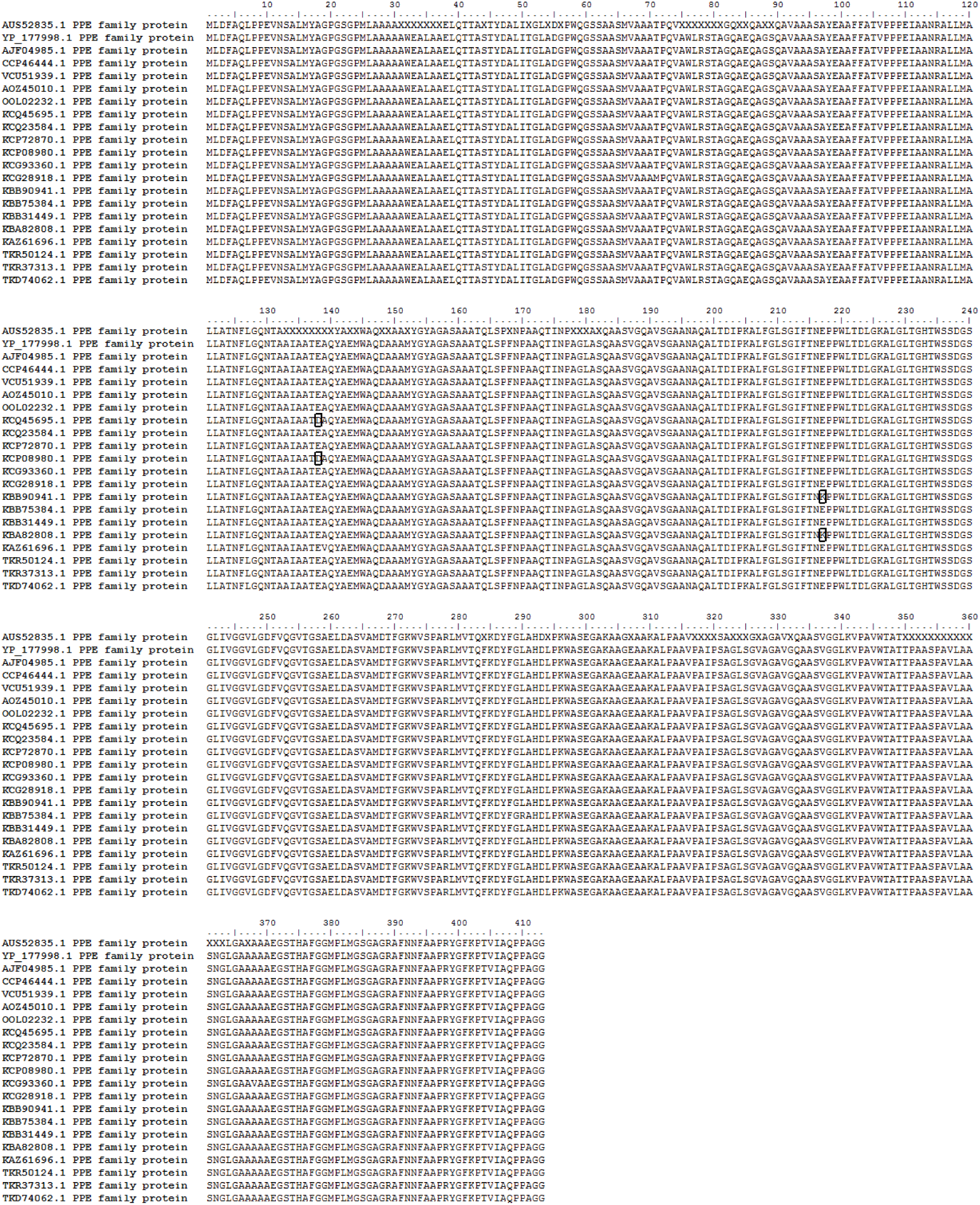
Multiple Sequence Alignment for partial sequence of the PPE65 protein in M. tuberculosis, using BioEdit software.

**Figure 2:**
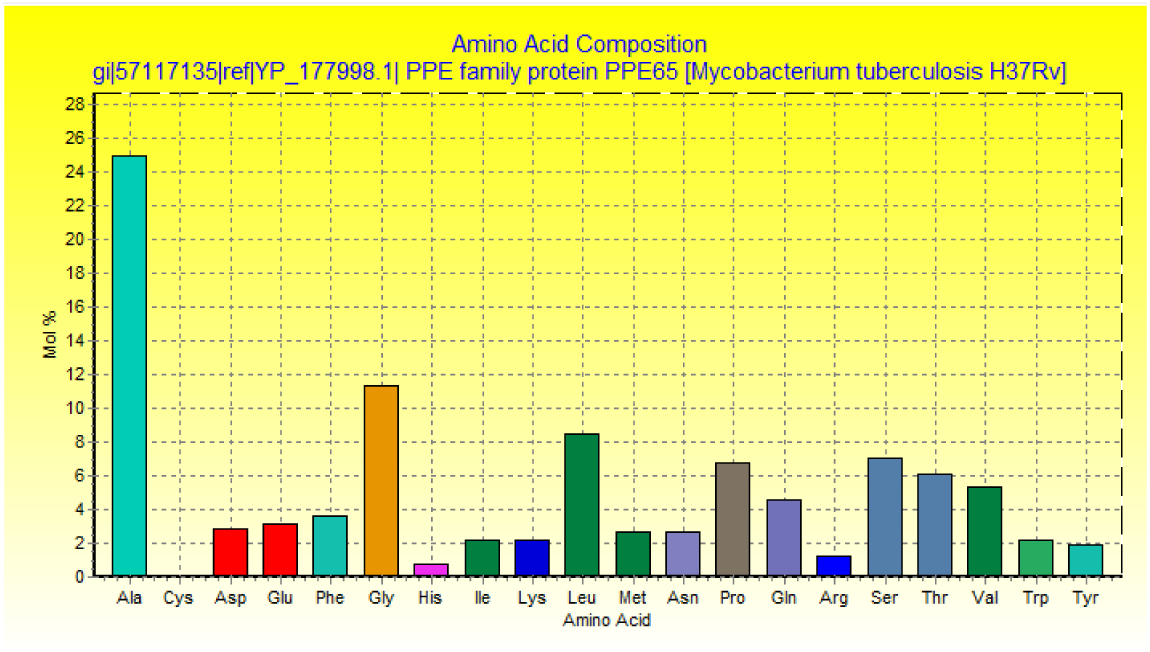
Aminoacid composition for *Mycobacterium Tuberculosis*PPE65 family protein using BioEdit software.

### B-cell epitope prediction

The reference sequence of *M. Tuberculosis* PPE65 was subjected to Bepipred linear epitope 2, Emini surface accessibility, Kolaskar & Tongaonkar antigenicity and Parker hydrophilicity prediction methods to test for various immunogenicity parameters (Table 2 and Figures 3-8). Two epitopes have successfully passed the three tests. Tertiary structure of the proposed B cell epitopes is illustrated (Figure 7 & 8).

**Table 2:** Conserved B cell epitopes that had successfully passed the tests.

| Peptide | Start | End | Length | Kolaskar & Tongaonkar antigenicity score (TH: 1.03) | Emini surface accessibility score (TH: 1) | Parker Hydrophilicity predictionscore (TH: 1.43) | ALLERTOP |
| --- | --- | --- | --- | --- | --- | --- | --- |
| YAGP | 17 | 20 | 4 | 1.041 | 1.24 | 2 | Non-allergen |
| QAAS | 183 | 186 | 4 | 1.039 | 1.212 | 4.175 | Non-allergen |
| QAAS | 333 | 336 | 4 | 1.039 | 1.212 | 4.175 | Non-allergen |

**Figure 3:**
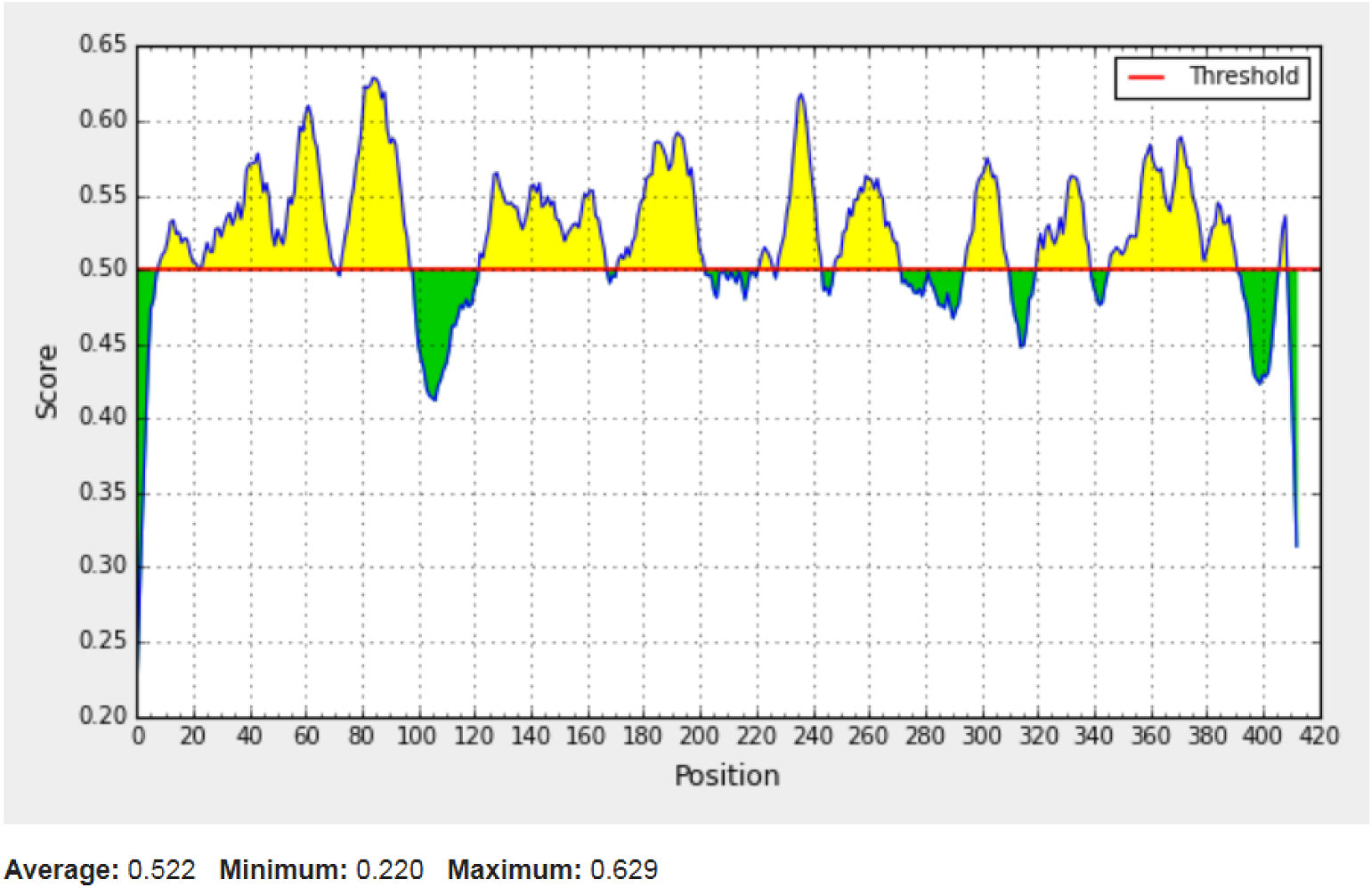
Bepipred Linear Epitope Prediction 2; Yellow areas above threshold (red line) are proposed to linearB-cell epitopes while the green areas are not.

**Figure 4:**
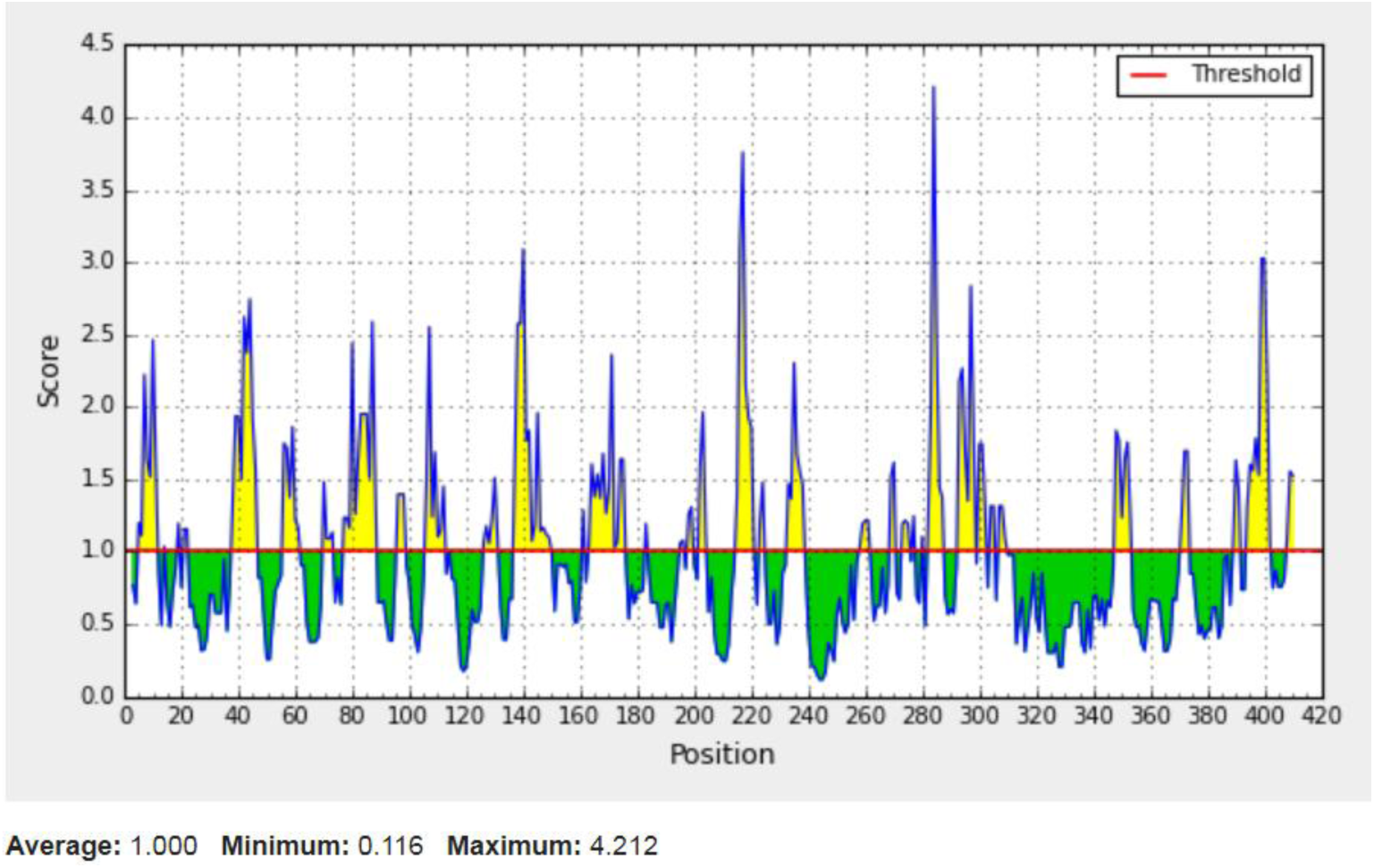
EMINI surface accessibility prediction; Yellow areas above the threshold (red line) are proposed to be a part of B cell epitopes and the green areas are not.

**Figure 5:**
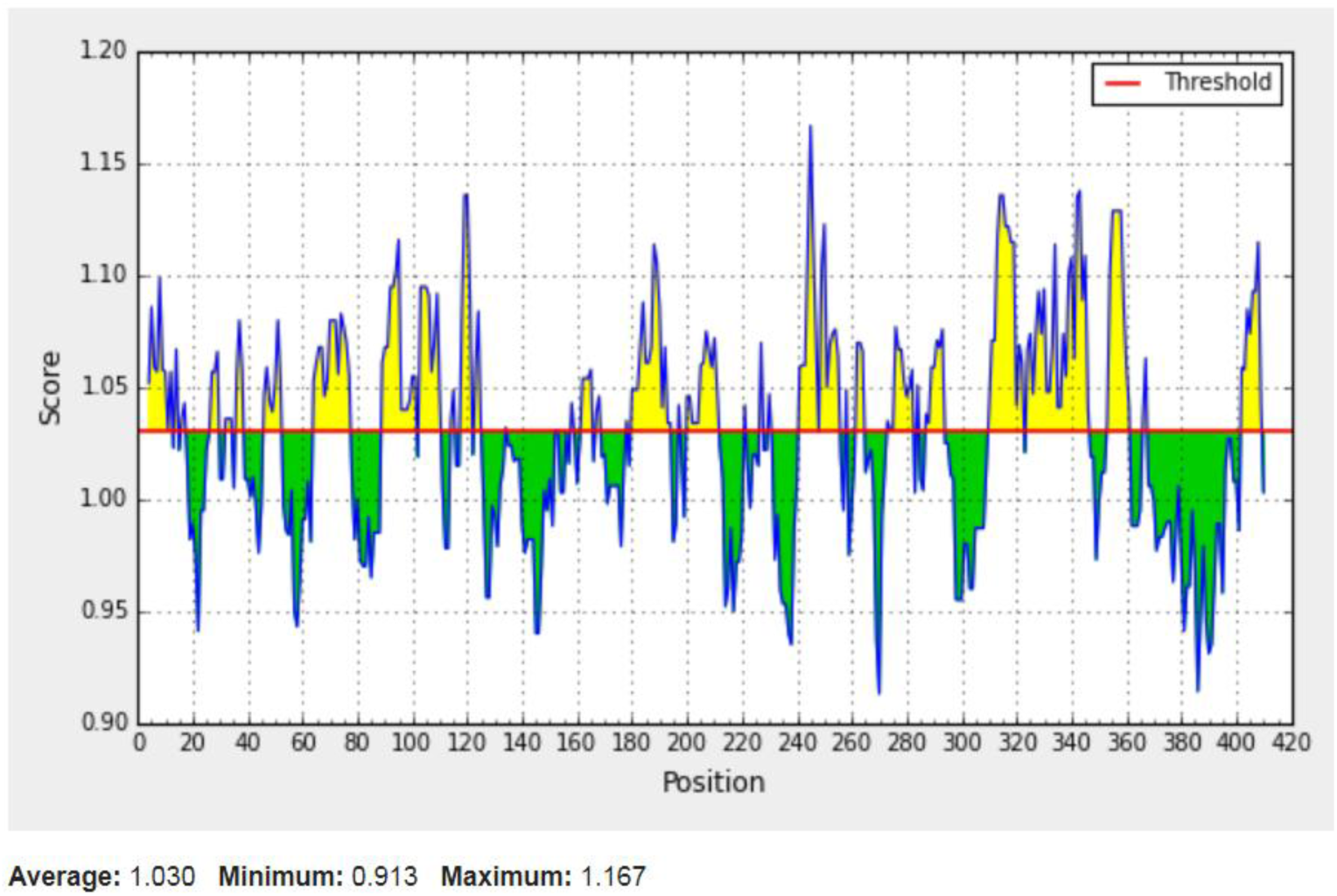
Kolaskar and Tonganokar antigenicity prediction; Yellow areas above the threshold (red line) are proposed to be antigenic B-cell epitopes and green areas are not.

**Figure 6:**
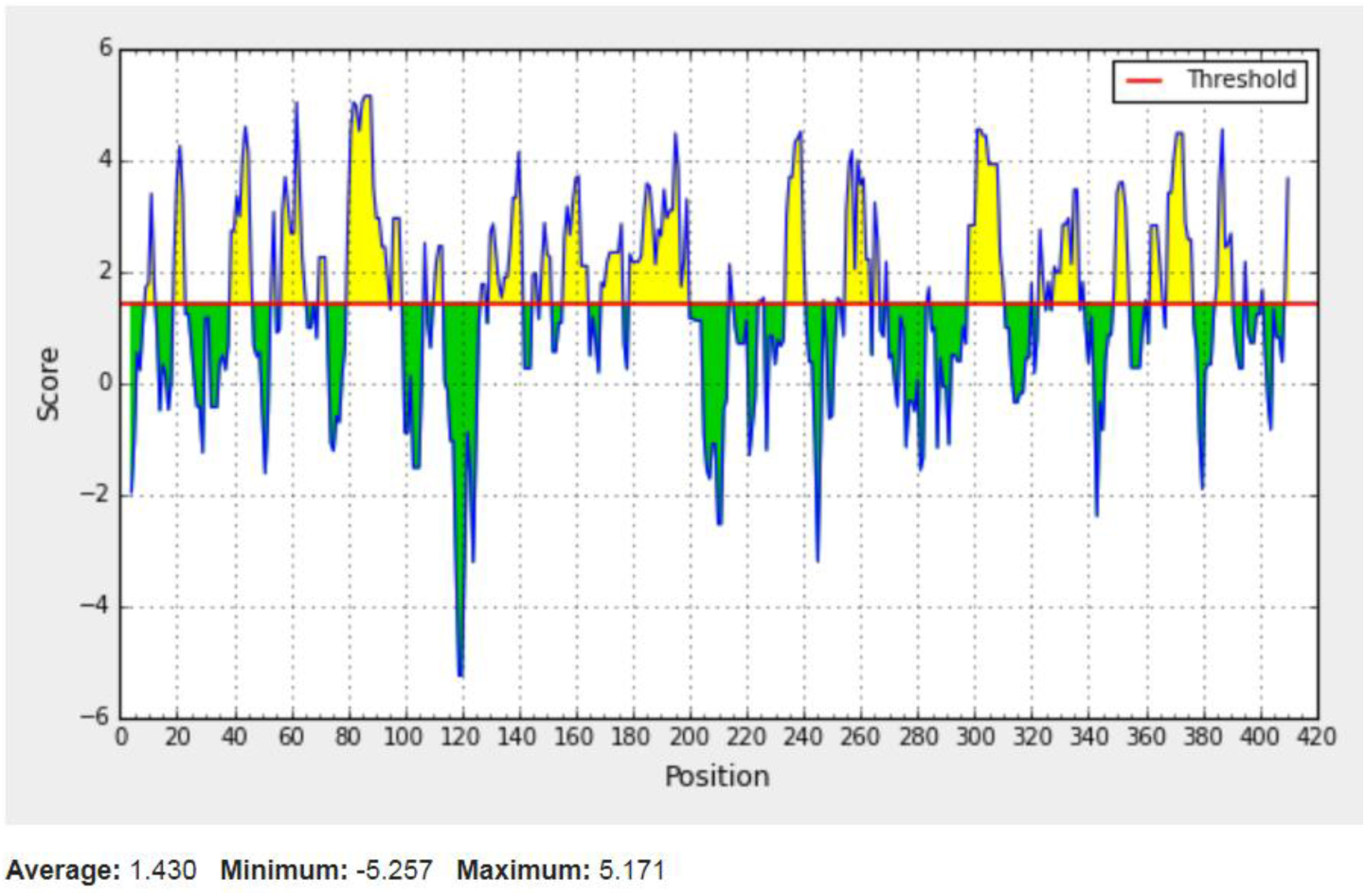
Parker Hydrophilicity prediction; Yellow areas above the threshold (red line) are proposed to be hydrophilic B-cell epitopes and green areas are not.

**Figure 7:**
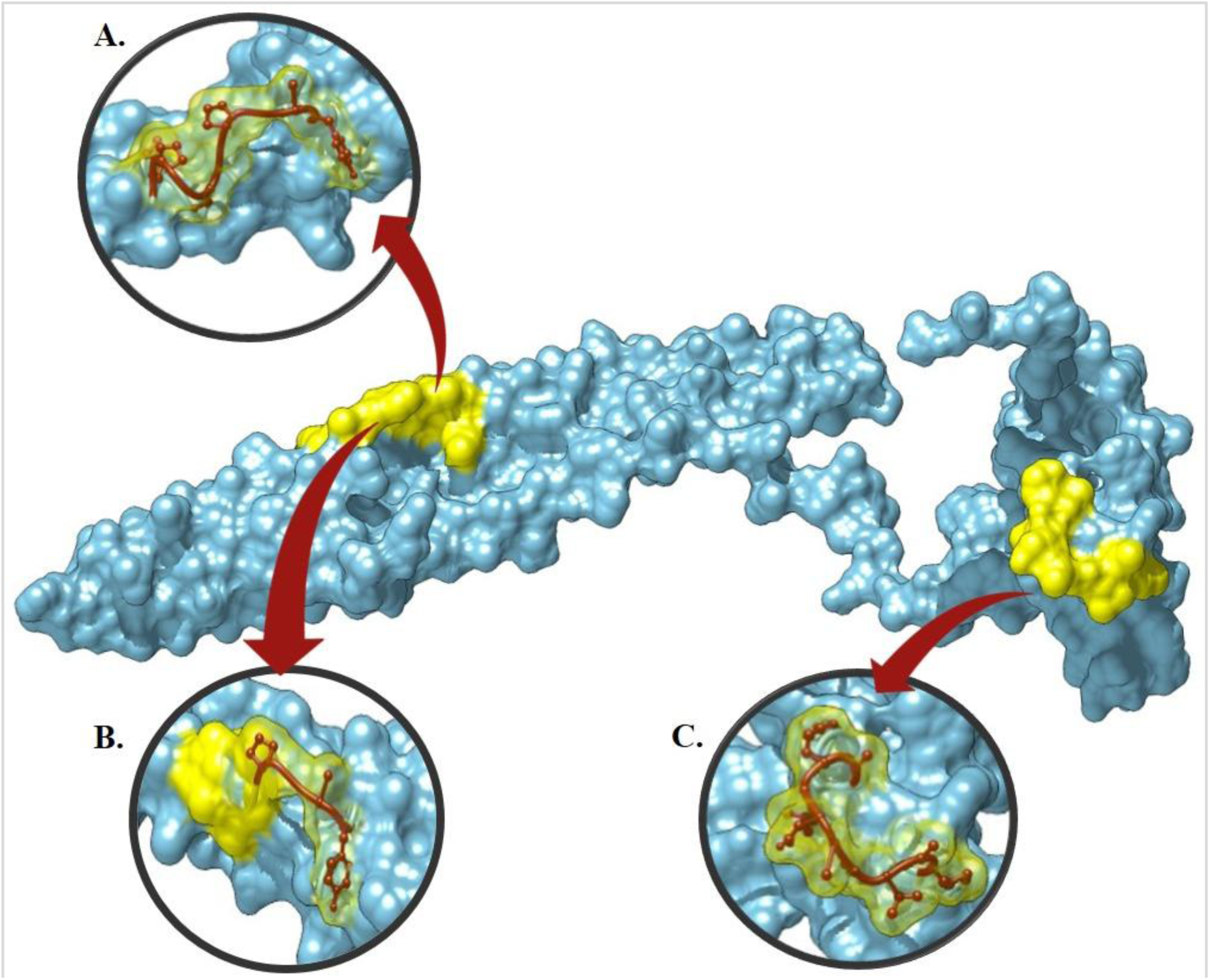
Three-dimensional visualisation of the most promising peptides in the study using Chimera (version 1.13.1rc) A. MHC-I binding T-cell peptide **YAGAPGSGPM** B. **YAGP** B-cell peptide which is a part of YAGPGSGPM and C. MHC-I binding T-cell peptide **AELDASVAM**.

### Prediction of Cytotoxic T-lymphocyte epitopes

The reference sequence was analyzed using (IEDB) MHC-1 binding prediction tool to predict T cell epitopes interacting with different types of MHC Class I alleles. 61 peptides were predicted to interact with different MHC-I alleles. The most promising epitopes with their corresponding MHC-1 alleles and IC50 scores are shown in (Table 3) followed by the tertiary structure of the candidate T cell epitope (Figure 7).

**Table 3:**
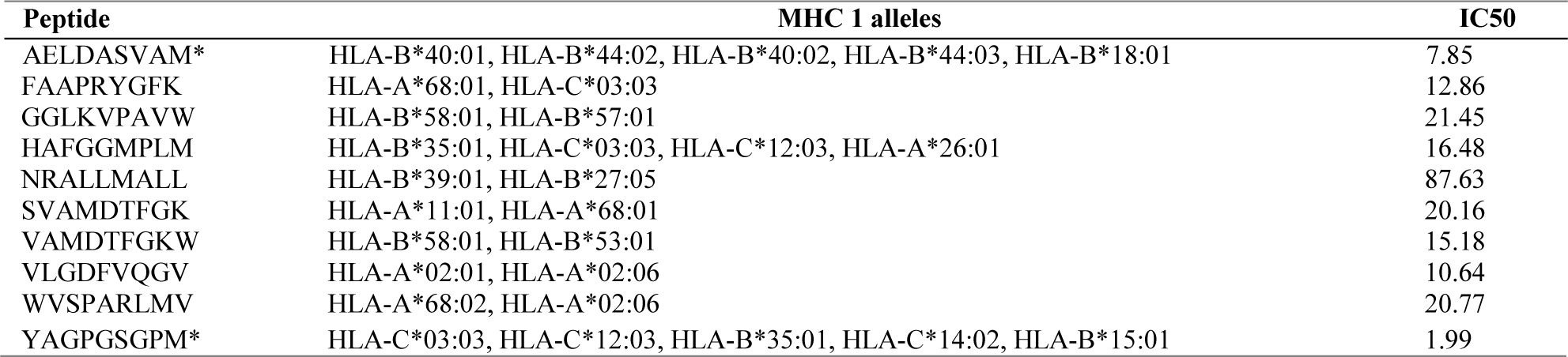
The most promising T cell epitopes and their corresponding MHC-1 alleles.

### Prediction of the Helper T-lymphocyte epitopes

Reference sequence was analysed using (IEDB) MHC-II binding prediction tool there were 85predicted epitopes found to interact with MHC-II alleles. The most promising epitopes with their corresponding alleles and IC50 scores are shown in (Table 4) along with the 3D structure of the proposed epitope (Figure 10).

**Table 4:**
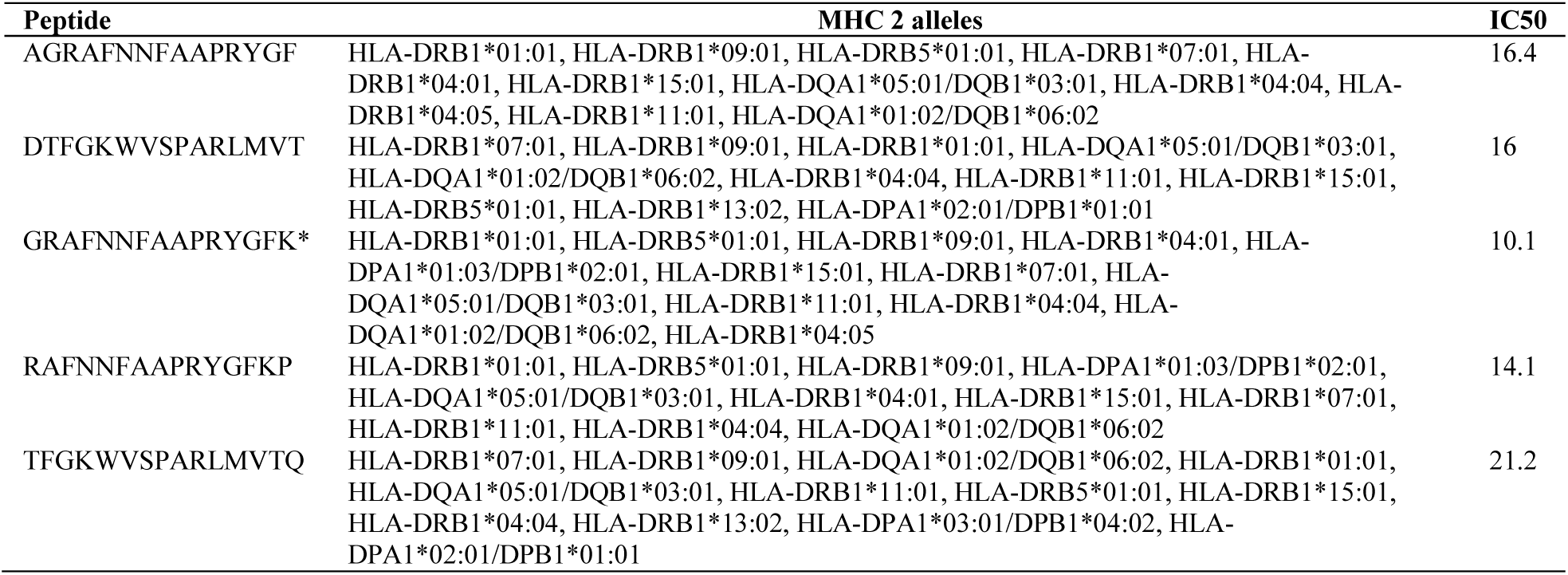
The most promising T cell epitopes and their corresponding MHC-2 alleles.

### Population Coverage Analysis

All MHC I and MHC II epitopes were assessed for population coverage against the whole world using IEDB population coverage tool. For MHC 1, epitopes with highest population coverage were VLGDFVQGV (40.6%) and YAGPGSGPM (33.81) (Figure 8 and Table 5). For MHC class II, the epitopes that showed highest population coverage were GRAFNNFAAPRYGFK (68.15%) and AGRAFNNFAAPRYGF (68.15%) (Figure 9 and Table 6). When combined together, the epitopes that showed highest population coverage were GRAFNNFAAPRYGFK (68.15) and AGRAFNNFAAPRYGF (68.15) (Figure10 and Table 7).

**Table 5:** Population coverage of proposed peptides interaction with MHC class I

| Epitope | Coverage Class 1 (%) | Total HLA hits |
| --- | --- | --- |
| VLGDFVQGV | 40.60% | 2 |
| YAGPGSGPM | 33.81% | 5 |
| AELDASVAM | 30.31% | 5 |
| HAFGGMPLM | 29.26% | 4 |
| RYGFKPTVI | 21.38% | 1 |
| SVAMDTFGK | 20.88% | 2 |
| FAAPRYGFK | 13.48% | 2 |
| APRYGFKPT | 12.78% | 1 |
| ALFGLSGIF | 8.44% | 1 |
| WQGSSAASM | 8.44% | 1 |

**Table 6:**
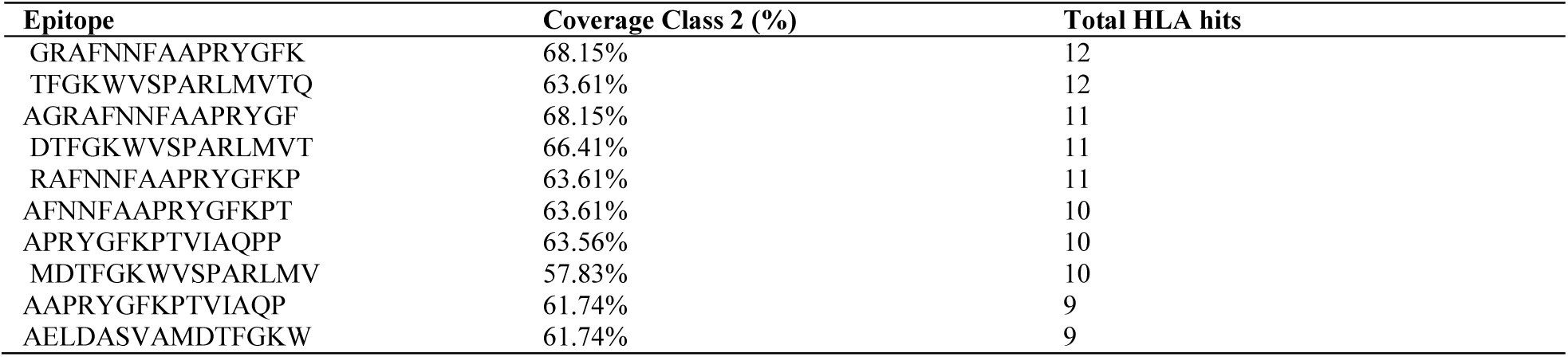
Population coverage of proposed peptides interaction with MHC class II

**Table 7:**
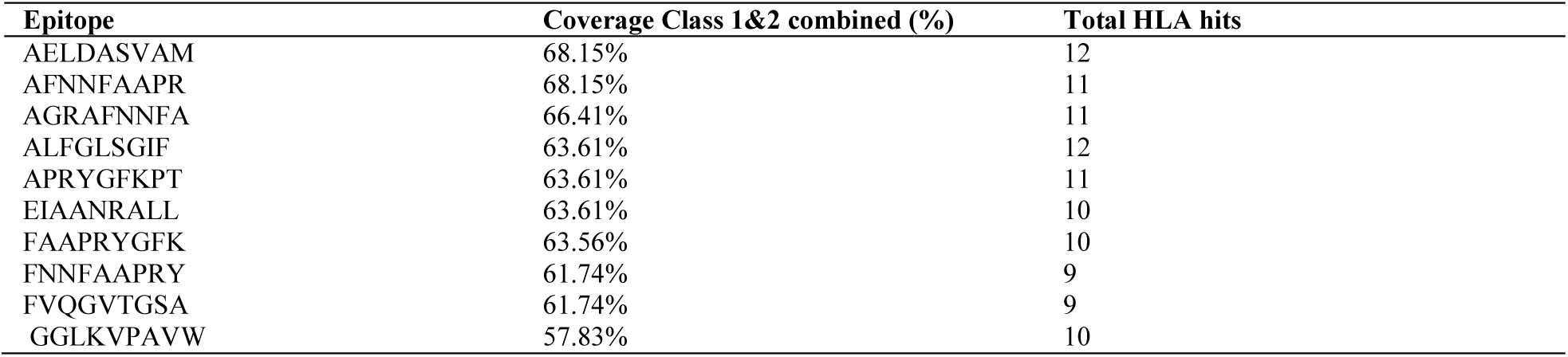
Population coverage of proposed peptide interaction with MHC class I&II combined

**Table 8:** The population coverage of whole world for the epitope set for MHC I, MHC II and MHC I&II combined.

| Country | MHC I | MHC II | MHC I&II combined |
| --- | --- | --- | --- |
| World | 95.34 % | 81.94 % * | 99.16% * |

**Figure 8:**
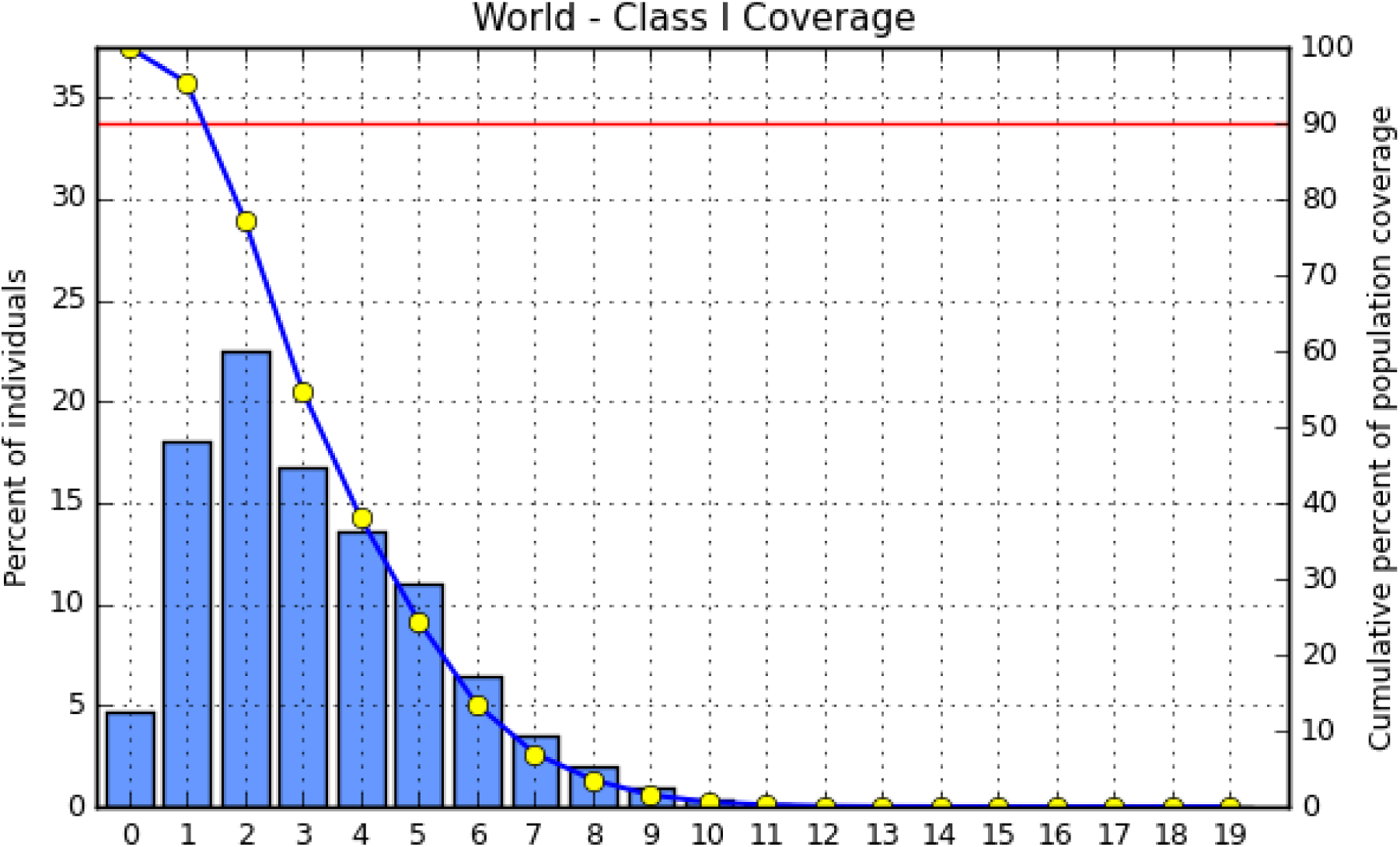
Population coverage for MHC class I epitopes.

**Figure 9:**
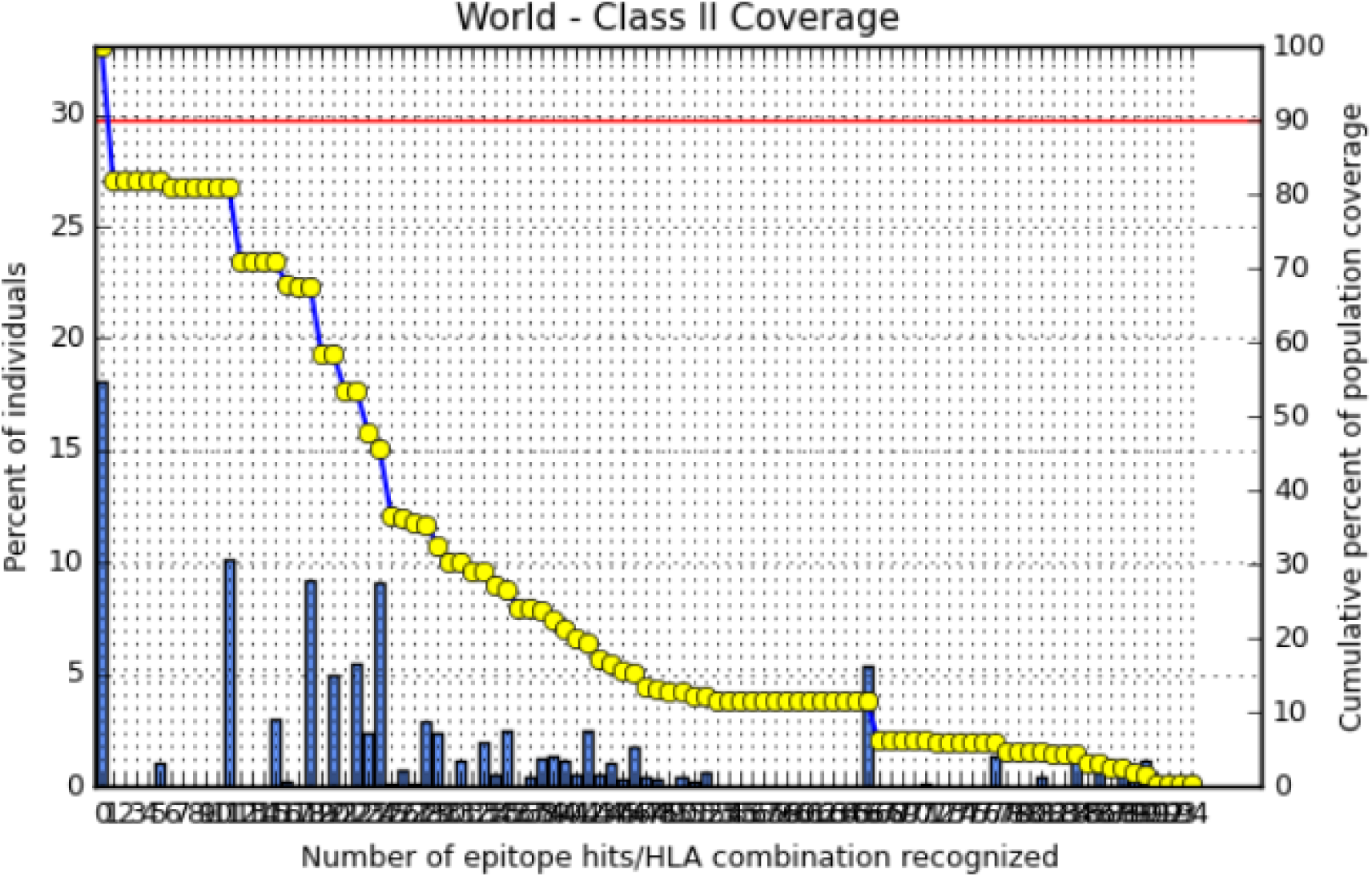
Population coverage for MHC class II epitopes.

**Figure 10:**
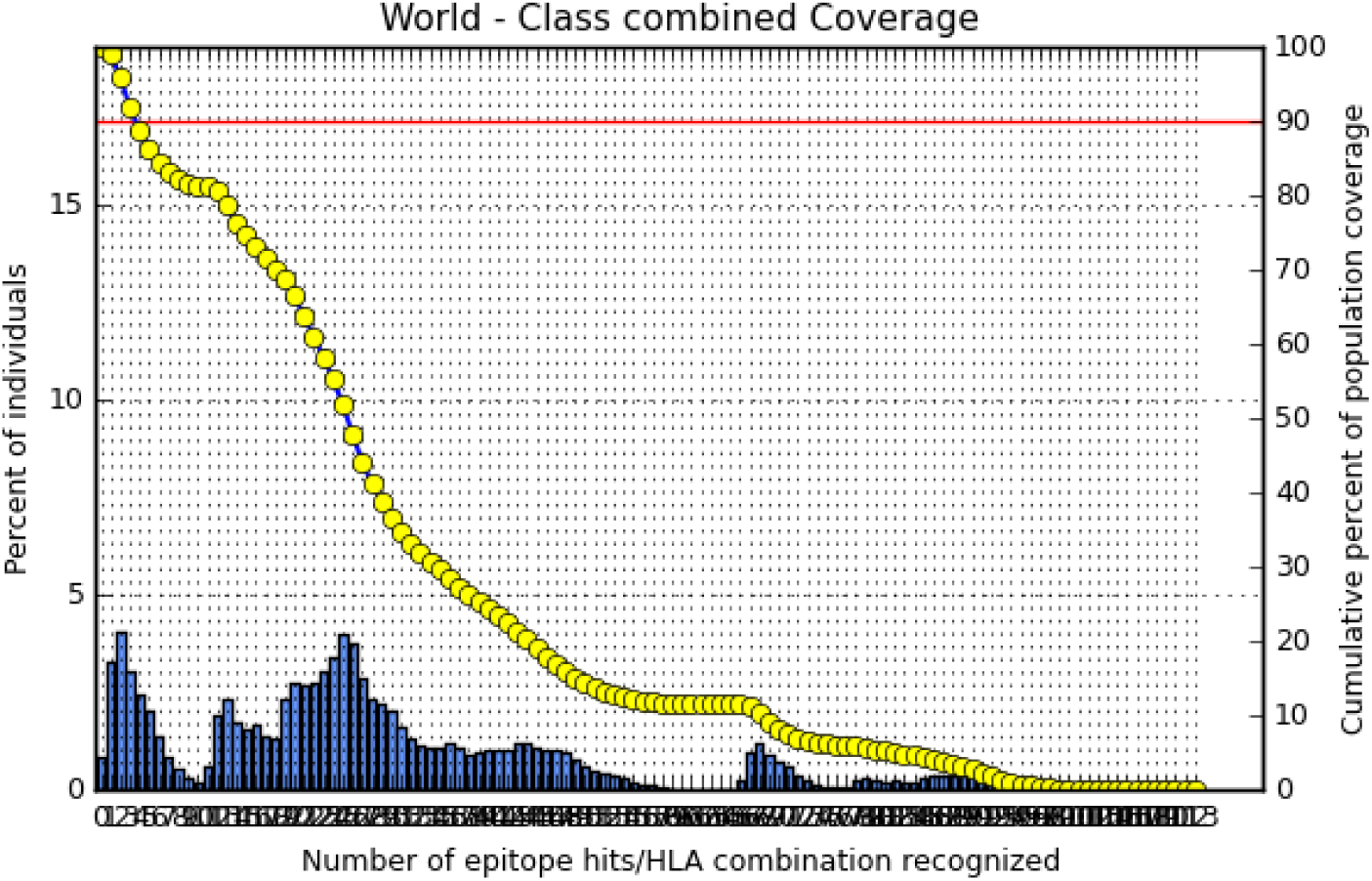
Population coverage for combined MHC I and II epitopes.

## DISCUSSION

This study suggests four most promising peptide candidates for a reverse vaccine design of M. tuberculosis targeting PPE65 protein, including three T-cell peptides (YAGPGSGPM, AELDASVAM, GRAFNNFAAPRYGFK) and a single B-cell peptide YAGP. These peptides make the strongest candidates due to their relatively high global population coverage, scoring the lowest rates of IC50 with their corresponding HLA alleles, indicating strong interaction between the peptide and allele.

This study also has a major focus on the analysis of T-cell peptides. Hence, tuberculosis is mainly maintained from dissemination by cell-mediated immunity.[67] CD4+ T cells are the principal antigen-specific cells responsible for containment of *M. tuberculosis* infection, although they can also be major contributors to disease during *M. tuberculosis* infection in several cases.[68]

For MHC-I, the length of peptides is strictly nine amino acids thus all peptides are nine amino acid long. Peptide YAGPGSGPM, is the most promising peptide in this study. It binds to five MHC-I alleles, (HLA-C*03:03, HLA-C*12:03, HLA-B*35:01, HLA-C*14:02, and HLA-B*15:01). Besides, scoring the lowest IC50 of 1.99 when binding to HLA-C*03:03 indicating very strong interaction. Moreover, the peptide’s global population coverage was the highest among the candidate peptides and was estimated for 33.81%. Another good finding which strengthens the point of selection of this particular peptide is that, a part of it YAGP peptide during the analysis of linear B-cell peptides was predicted to elicit a good immune response according to antigenicity, surface accessibility, hydrophilicity and linearity tests. This indicates that YAGPGSGPM is at least partially located on the surface of the protein and that it’s immunogenic stimulating both B and T-cell responses.

Peptide AELDASVAM is also a strong possible peptide for MHC-I related immune responses. Not only does the peptide bind to five HLA alleles (HLA-B*40:01, HLA-B*44:02, HLA-B*40:02, HLA-B*44:03, HLA-B*18:01), but it has a very low IC50 of 7.85 to HLA-B*40:01 allele. The peptide is predicted to cover 30.31% of the total global population.

As for MHC-II alleles, the proposed alleles were longer than nine peptides to give more space for manipulation during invivo and in vitro analysis. GRAFNNFAAPRYGFK peptide is the most promising peptide among the suggested peptides. It binds to a vast number of MHC-II alleles, approximately 12 alleles. In addition, it scored lowest IC50 of 10.1 to HLA-DRB1*01:01and the highest population coverage with a score of 68.15%.

In this study, twenty-three peptides were shared in eliciting both MHC-I and MHC-II immune responses. Among these peptides were YAGPGSGPM, and AELDASVAM which proves the remarkable efficiency of the selected epitopes. This is advantageous when designing a peptide-based vaccine as it reduces the vast number of selected epitopes.

Different peptide based vaccine approaches for TB have been reported previously. A study by Rai PK et al. 2017 has reported that a lapidated peptide was introduced (L91) and engaged in an animal model trails to detect the efficiency of the peptide in activating T-helper cells. The result then confirmed that L91 peptide led to enhanced immune response to BCG vaccine.[69] A study by Coppola et al. 2015, suggests that a long synthetic peptide derived from Latency antigen Rv1733c of M. tuberculosis which is highly expressed in patients with dormant TB, can efficiently elicit an immune response, particularly T-cell response. The result was validated by using animal models for in vivo response.[70]

In this study, and besides being strictly computational, another limitation was found in population coverage analysis of MHC II; a total of twelve alleles did not give predictions. This includes (HLA-DQA1*05:01/DQB1*03:01, HLA-DQA1*01:02/DQB1*06:02, HLA-DPA1*01/DPB1*04:01, HLA-DPA1*01:03/DPB1*02:01, HLA-DQA1*03:01/DQB1*03:02, HLA-DRB3*01:01, HLA-DRB4*01:01, HLA-DRB5*01:01, HLA-DQA1*05:01/DQB1*02:01, HLA-DPA1*03:01/DPB1*04:02, HLA-DQA1*04:01/DQB1*04:02, and HLA-DPA1*02:01/DPB1*01:01). In addition, the tertiary structure of the PPE65 protein was incomplete and this resulted in failure of visualization of the promising MHC-II peptide GRAFNNFAAPRYGFK. We recommend future researches conducting an in vivo and in vitro analysis of the proposed peptides and a population coverage analysis of the HLA alleles that did not give result along with homology modelling for PPE65 protein.

Peptide vaccines are an attractive alternative to conventional vaccines, as their concept relies on usage of short peptide fragments to induce a highly targeted immune responses, consequently avoiding allergenic and/or reactogenic sequences. [44]Reverse vaccine technology has proved its efficiency as many of the designed vaccines have made it into clinical trials. For example, a multiple peptides vaccine derived from tumor-associated antigens (TAAs) made it to phase II clinical trial for head and neck squamous cell cancer (HNSCC). A phase I trial was registered with University Hospital Medical Information Network (UMIN) number UMIN000002022, with a fixed 2-mg dose of KIF20A and VEGFR1 peptides for patients with advanced unresectable pancreatic cancer targeting nineteen patients sub grouped into the HLA-A*2402-positive group and HLA-A*2402-negative group respectively. Results of this study showed that vaccination with KIF20A and VEGFR1 peptides was a safe treatment, and might also be a promising treatment as HLA-A*2402-positive group had an increased survival rate in comparison to the negative group.[71]

In general, peptide based vaccines are safer, more stable with a minimal exposure to hazardous or allergenic substances compared to conventional vaccines. Moreover, these peptide vaccines are less labouring, time consuming and cost efficient. The only weakness regarding them is the need to introduce an adjuvant to increase immunogenic stimulation of the vaccine recipient. [44][72][73][74]

## CONCLUSION

Since the need for a proper vaccine for M. tuberculosis is increasing with time, along with the reduced response to BCG vaccine, and growing population rates, poverty and pollution. This study suggests four peptides suitable for further analysis using reverse vaccinology techniques. YAGPGSGPM, AELDASVAM, and GRAFNNFAAPRYGFK peptides have acceptable global population coverage percentages, good binding affinity to MHC alleles.

